# Cyclic Nucleotide-Gated Ion Channel 2 modulates auxin homeostasis and signaling

**DOI:** 10.1101/508572

**Authors:** Sonhita Chakraborty, Masatsugu Toyota, Wolfgang Moeder, Kimberley Chin, Alex Fortuna, Marc Champigny, Steffen Vanneste, Simon Gilroy, Tom Beeckman, Eiji Nambara, Keiko Yoshioka

## Abstract

Cyclic Nucleotide Gated Ion Channels (CNGCs) have been firmly established as Ca^2+^-conducting ion channels that regulate a wide variety of physiological responses in plants. CNGC2 has been implicated in plant immunity and Ca^2+^ signaling due to the autoimmune phenotypes exhibited by null mutants of *CNGC2*. However, *cngc2* mutants display additional phenotypes that are unique among autoimmune mutants, suggesting that CNGC2 has functions beyond defense and generates distinct Ca^2+^ signals in response to different triggers. In this study we found that *cngc2* mutants showed reduced gravitropism, consistent with a defect in auxin signaling. This was mirrored in the diminished auxin response detected by the auxin reporters DR5::GUS and DII-VENUS and in a strongly impaired auxin-induced Ca^2+^ response. Moreover, the *cngc2* mutant exhibits higher levels of the endogenous auxin indole-3-acetic acid (IAA), indicating that excess auxin in *cngc2* causes its pleiotropic phenotypes. These auxin signaling defects and the autoimmunity syndrome of *cngc2* could be suppressed by loss-of-function mutations in the auxin biosynthesis gene *YUCCA6* (*YUC6*), as determined by identification of the *cngc2* suppressor mutant *repressor of cngc2* (*rdd1*) as an allele of *YUC6*. A loss-of-function mutation in the upstream auxin biosynthesis gene *TRYPTOPHAN AMINOTRANSFERASE OF ARABIDOPSIS* (*TAA1, WEAK ETHYLENE INSENSITIVE8*) also suppressed the *cngc2* phenotypes, further supporting the tight relationship between CNGC2 and the TAA–YUC-dependent auxin biosynthesis pathway. Taking these results together, we propose that the Ca^2+^ signal generated by CNGC2 is a part of the negative feedback regulation of auxin homeostasis in which CNGC2 balances cellular auxin perception by influencing auxin biosynthesis.

## INTRODUCTION

Calcium (Ca^2+^) is a ubiquitous second messenger that orchestrates many signaling pathways in eukaryotes. A diverse range of stimuli elicit transient changes in the cytosolic free Ca^2+^ concentration ([Ca^2+^]_cyt_) in plants. These include developmental cues, abiotic stresses such as drought, heat, and wounding, and biotic stimuli such as interactions with pathogenic and symbiotic microorganisms (Moeder et al., 2019; Yuan et al., 2017). Each type of stimulus is thought to generate a distinctive spatio-temporal ‘Ca^2+^ signature’, which is decoded by the direct binding of Ca^2+^ to calcium sensor proteins such as calmodulins (CaM), Calcium-dependent protein kinases (CDPKs), and others (Edel et al., 2017). These sensor proteins may undergo conformational changes upon binding and interact with or phosphorylate various target substrates to regulate downstream factors (DeFalco et al., 2010; Edel et al., 2017). The Ca^2+^ flux is regulated by a combination of various channels, pumps, and transporters, which move Ca^2+^ to and from extracellular and/or intracellular Ca^2+^ stores to the cytoplasm (Demidchik et al., 2018). Despite the central role played by Ca^2+^ in plant physiological responses, the identity of plant Ca^2+^ channels that regulate specific [Ca^2+^]_cyt_ remains elusive.

Cyclic nucleotide gated ion channels (CNGCs) function as Ca^2+^ channels and are linked to Ca^2+^ signaling in plants (DeFalco et al., 2016; Dietrich et al, 2020; Jammes et al., 2011; Zelman et al., 2012, Moeder et al., 2019). They are involved in a variety of physiological processes that are regulated by Ca^2+^ signaling, such as pollen tube growth, thermo-sensing, pathogen resistance, root growth, and symbiotic interactions (Brost et al., 2019; Charpentier et al., 2016; Finka et al., 2012; Moeder et al., 2011; Dietrich et al, 2020). The involvement of CNGCs in plant defense was first suggested in a study of the Arabidopsis null mutant of *CNGC2, <u>d</u>efense, <u>n</u>o <u>d</u>eath<u>1</u>* (*dnd1, also known as cngc2-1*). *cngc2* mutants exhibit autoimmune phenotypes such as stunted growth, conditional spontaneous cell death, and elevated basal levels of salicylic acid (SA), that confer enhanced resistance to various pathogens (Clough et al., 2000; Yu et al., 1998). They have a reduced ability to mount a hypersensitive response (HR), which is a form of programmed cell death (PCD) around the site of pathogen infection, often observed during effector triggered immunity (Yu et al., 1998). Two null mutants of *CNGC4, <u>H</u>R-like lesion <u>m</u>imic<u>1</u>* (*hlm1*; Balagué et al., 2003) and *<u>d</u>efense, <u>n</u>o <u>d</u>eath<u>2</u>* (*dnd2*, later referred to as *cngc4*; Jurkowski et al., 2004), and a gain-of-function mutant of *CNGC12, <u>c</u>onstitutive expresser of <u>p</u>athogenesis-related genes<u>22</u>* (*cpr22*), also show alterations in defense responses (Yoshioka et al., 2006). The inhibition of HR-like spontaneous cell death in *cpr22* by Ca^2+^ channel blockers, and the higher [Ca^2+^]_cyt_ levels in *cpr22* further support the notion that *CNGC12* induces defense response by activating Ca^2+^ signalling (Urquhart et al., 2007; Moeder et al., 2011, 2019;). In contrast, the *cngc2* mutant displays a reduced HR phenotype and suppression of Ca^2+^ signals induced by pathogen elicitors, suggesting that CNGC2 positively regulates defense (Ali et al., 2007, Tian et al., 2019). However, the autoimmune phenotype of *cngc2* contradicts this notion (Moeder et al., 2011; Dietrich et al., 2020).

The *cngc2* mutant is hypersensitive to elevated Ca^2+^ levels (Chan et al., 2003). A genome-wide transcriptional study revealed that the gene expression pattern of *cngc2* resembles that of wild type under elevated Ca^2+^ stress (Chan et al., 2008). This observation was supported by a study reporting that CNGC2 maintains Ca^2+^ homeostasis by facilitating Ca^2+^ unloading from the vasculature into leaf cells (Wang et al., 2017). The Ca^2+^ hypersensitivity of *cngc2* may be the cause of the autoimmune phenotypes (high SA levels, H_2_O_2_ accumulation, and cell death), since these are largely suppressed when *cngc2* seedlings are grown in media with low [Ca^2+^] (Chan et al., 2003; Wang et al., 2017; Tian et al., 2019). *cngc2* mutants are impaired in pathogen-associated molecular pattern (PAMP)-induced immunity (PTI) under Ca^2+^ concentrations that induce the pleiotropic phenotypes of this mutant, suggesting a complex relationship between Ca^2+^ concentration and immunity in this mutant (Tian et al., 2019).

In addition to these immunity phenotypes, multiple studies reported roles of CNGC2 in abiotic stress responses and development. For example, *cngc2* plants display delayed flowering. This is not typical for conventional SA-accumulating autoimmune mutants, which usually exhibit early flowering (Chin et al., 2013; Fortuna et al., 2015). This finding suggests that the autoimmune phenotype observed in *cngc2* may not be simply due to elevated SA levels. CNGC2 has also been suggested to play a role in CLAVATA3/CLAVATA1 (CLV3/CLV1) signaling in shoot apical meristem (SAM) maintenance (Chou et al., 2016). CLV1, a plasma membrane-localized receptor kinase, and its peptide ligand CLV3 regulate cell differentiation (Somssich et al., 2016). Most recently CNGC2 is also implicated in light stress induced Ca^2+^ signaling (Fichman et al., 2021). Jointly, these observations indicate that CNGC2 could be activated by a diverse set of signals, feeding into immunity, stress tolerance, and developmental outputs (Dietrich et al, 2020).

The plant hormone auxin plays a central role throughout plant development and its activity is typically associated with local accumulation that triggers a developmental response (Vanneste & Friml, 2009). The most bioactive endogenous auxin, indole-3-acetic acid (IAA), is produced from tryptophan via indole-3-pyruvate (IPA) in a two-step reaction, involving the TRYPTOPHAN AMINOTRANSFERASE OF ARABIDOPSIS (TAA) and YUCCA (YUC) enzyme families in Arabidopsis (Mashiguchi et al., 2011; Stepanova et al., 2011; Won et al., 2011). The YUC proteins belong to the flavin-containing monooxygenase (FMO) enzyme family and play important roles in growth and development via auxin production (Cheng et al., 2006). For example, the *yuc* mutants and *YUC* over-expression lines show defects in pollen development, embryogenesis, senescence process and leaf morphology (Cao et al., 2019; Cheng et al., 2007; Kim et al., 2011). In addition, *YUC* genes are involved in stress responses such as heat and drought stress as many environmental stimuli converge on this pathway to modify plant growth and development (Cao et al., 2019).

To study CNGC2-mediated signal transduction, we screened for suppressors of a *cngc2* null mutant and identified *repressor of <u>d</u>efense, no <u>d</u>eath<u>1</u>* (*rdd1*; Chin et al., 2013). The *rdd1* mutation supresses almost all *cngc2*-related phenotypes, except its Ca^2+^ hypersensitivity, indicating that *RDD1* acts downstream of CNGC2’s channel function. In this study, we identified the causal mutation of *rdd1* as a loss-of-function mutation in the auxin biosynthesis gene, *YUCCA6* (*YUC6*). Here, we propose that CNGC2 plays a role in auxin homeostasis in addition to immunity as *cngc2* is defective in canonical auxin signaling and auxin-induced Ca^2+^ signaling, phenotypes that can also be all explained by hyper-accumulation of endogenous IAA.

## RESULTS

### *rdd1* is a loss-of-function allele of *YUC6*

The CNGC2 null mutant *dnd1* results from a point mutation. In this study as well as Chin et al (2013), we have used the *cngc2-3* T-DNA insertion knockout line (a *CNGC2* T-DNA insertion line in the Columbia background). To clone the causal locus for *rdd1, rdd1 cngc2-3* plants were outcrossed with *cngc2-1* (in the Wassilewskija [Ws] background) to generate a mapping population. Using 705 plants of this F_2_ mapping population, the causal locus for *rdd1* was mapped to an approximately 800-kb region that contained 193 coding sequences (Chin et al., 2013). Using whole-genome sequencing, the causal mutation of *rdd1* was narrowed down to four candidate genes within this region: *AT5G24680, AT5G25590, AT5G25620*, and *AT5G26050*. Of these, only the mutation in *AT5G25620* was located in a coding region. Using qRT-PCR analysis, we found that *rdd1* did not have significant alterations in the expression of the three other candidate genes **(Supplemental Fig. S1)**. Therefore, the *rdd1* mutation is likely a non-synonymous amino acid change from proline to leucine at residue 297 in the third exon of *AT5G25620*, which encodes the protein YUC6, a flavin-containing monooxygenase-like (FMO) protein involved in a key step in auxin biosynthesis (**Fig. 1A**; Mashiguchi et al., 2011; Won et al., 2011).

**Figure 1.**
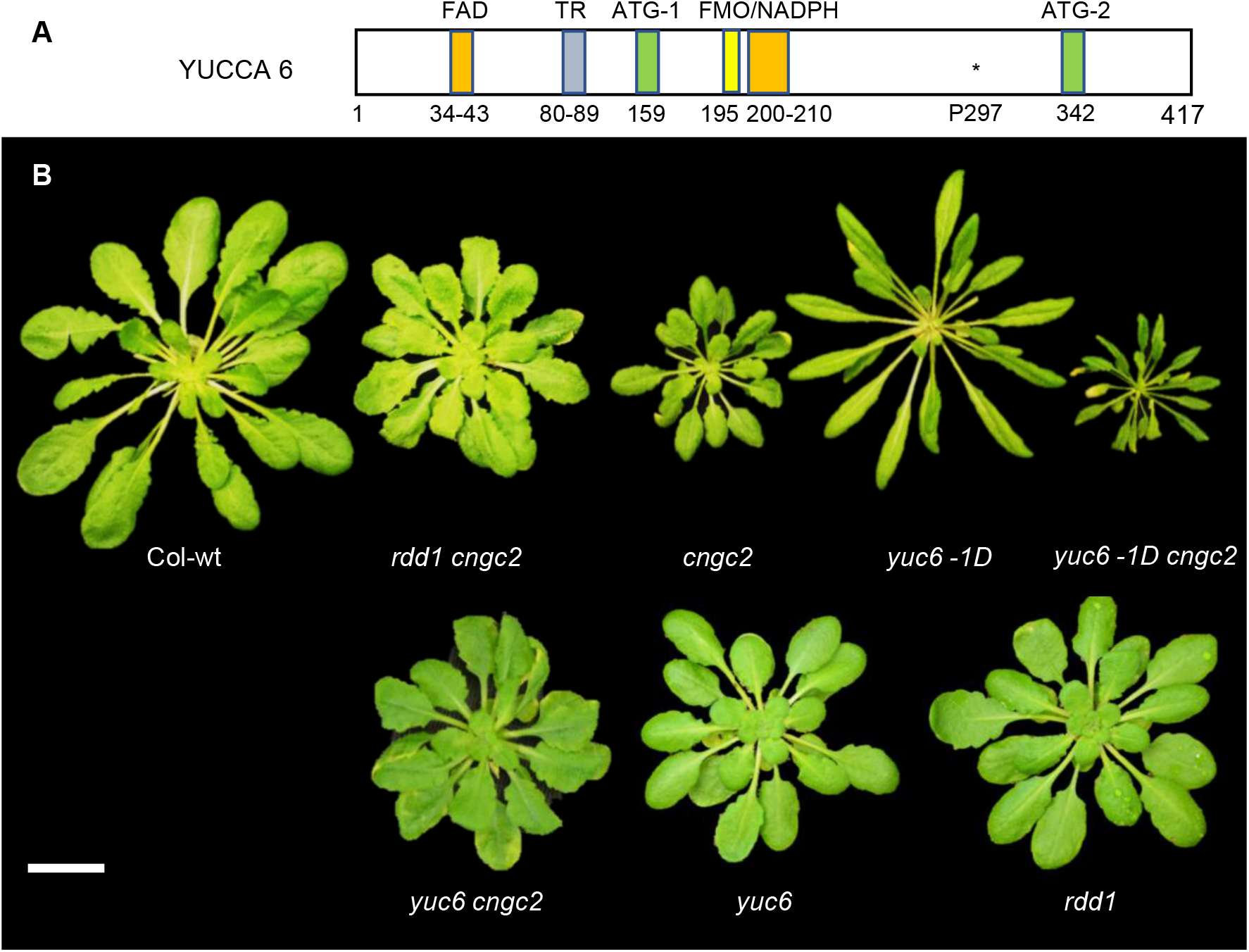
*rdd1* is a loss-of-function allele of *YUCCA6*. A, Schematic representation of the *rdd1* mutation and catalytic sites in YUCCA6 (YUC6). The relative locations of the FAD-binding domain (FAD, a.a. 34–43), thiol reductase domain (TR, a.a. 80-89), FMO identity sequence (FMO, a.a.195-199), and NADPH-binding domain (NADP, a.a. 202–210), the ATG-containing motifs (ATG1, a.a 159; and ATG2, a.a. 342), and the *rdd1* mutation position P297 are depicted based on gene model YUC6.1 TAIR). B, Morphology of approximately 5-week-old plants grown in short day conditions (9L:15D). *yuc6 cngc2* partially suppress *cngc2*-conferred dwarf morphology to the same degree as *rdd1 cngc2. yuc6*, the YUC6 knockout line, has shorter and wider rosette leaves compared to Col-wt, similar to the *rdd1-1* single mutant. *yuc6-1D* is unable to suppress *cngc2*-conferred dwarf phenotypes. Yellowing of leaf tip is observed in *yuc6-1D cngc2*. Scale bar = 1 cm.

*rdd1* is a point mutation that does not cause a premature stop codon or frame shift in the *YUC6* gene. Furthermore, the *rdd1* mutation was not located in or in proximity of any of the known domains of YUC6 (Fig. 1A); thus, its suppression of the *cngc2* phenotype could be due to either a gain- or loss-of-function of this enzyme. To determine which of these was the case, we tested *yuc6-1D*, a *YUC6* overexpressing activation line (Kim et al., 2007), and *yucca6-3k* (henceforth, *yuc6*), a *YUC6* knockout mutant (Cheng et al, 2006), for their ability to suppress *cngc2* phenotypes. First, we introduced the *yuc6-1D* allele into *cngc2*. None of the 132 F_2_ generation plants analyzed, resulting from a cross between *yuc6-1D* and *cngc2* showed a *rdd1*-like morphological phenotype. Instead, we observed seedlings that had a smaller stature than either parental line (enhanced dwarfism), while displaying a typical *yuc6-1D* phenotype (such as long, narrow leaves with elongated petioles; **Fig. 1B**, Kim et al., 2007). Thus, we genotyped the *yuc6-1D* and *cngc2* status of 47 F_2_ progenies (**Supplemental Table S1**). We found that these smaller plants were homozygous for *cngc2* and *yuc6-1D* positive. Therefore, *yuc6-1D* does not rescue the dwarf phenotype of *cngc2*; rather, it enhances this phenotype (Indicated as ss in **Supplemental Table S1**).

By contrast, the loss-of-function mutant *yuc6* suppressed the *cngc2* phenotype. Indeed, the *yuc6 cngc2* homozygotes in an F_2_ population derived from a cross between *yuc6* and *cngc2* were identical in phenotype to *rdd1 cngc2* **(Fig. 1B)**, indicating that *rdd1* is a loss-of-function allele of *YUC6*. In addition, the *rdd1* single mutant exhibited the identical broad leaf phenotype as *yuc6*, further supporting the notion that *rdd1* is a loss-of-function allele of *YUC6* **(Fig. 1B)**. The *cngc2*-like plants were more numerous in this F_2_ population than would be expected if *rdd1* (and therefore also *yuc6)* were dominant in its suppression of *cngc2* (Chin et al., 2013). Therefore, we conducted additional detailed genetic analyses using two backcrossed populations of *yuc6 cngc2* x *cngc2* and *rdd1 cngc2* x *cngc2*. We found that *rdd1* is a semi-dominant mutation with a dosage effect (**Supplemental Table S2, S3**, and **Supplemental Fig. S2**). The difference from the previously published *rdd1* segregation analysis (Chin et al., 2013) is probably due to the environmental sensitivity of the penetrance of *cngc2* phenotypes. Taken together, these results show that *rdd1* is a loss-of-function mutant of *YUC6* and that the *cngc2* phenotype depends on the dose of YUC6.

### *cngc2* phenotypes depend on functional *YUC6*

The *cngc2* mutant phenotype is complex as it displays a range of seemingly unrelated phenotypes, all of which are suppressed in *rdd1 cngc2*, with the exception of Ca^2+^ hypersensitivity (Chin et al., 2013). Therefore, we analyzed the YUC6-dependence of the *cngc2* phenotypes. As seen in *rdd1 cngc2* (Chin et al. 2013), trypan blue staining of four-week-old plants revealed that *yuc6 cngc2* plants exhibited less spontaneous cell death than *cngc2* plants **(Fig. 2A)**. Moreover, *yuc6 cngc2* plants partially lost the enhanced resistance of *cngc2* to the oomycete pathogen *Hyaloperonospora arabidopsidis* (*Hpa*) isolate Noco2 **(Fig. 2B** and **C)**. Accordingly, increased basal levels of SA in *cngc2* was suppressed to the same degree in *yuc6 cngc2* as in *rdd1 cngc2* (**Fig. 2D**). We have also analyzed the basal SA levels of the *yuc6* single mutants multiple times and did not find a statistically significant alteration in these mutants, indicating that *yuc6* and *rdd1* epistatically suppress increased SA levels in *cngc2* (**Supplemental Fig. S3**). Furthermore, *cngc2* exhibits delayed flowering, which is suppressed in *rdd1 cngc2* in an SA-independent manner (Fortuna et al., 2015). Like *rdd1 cngc2*, the *yuc6 cngc2* plants showed suppression of the delayed flowering and even displayed earlier flowering, compared to WT **(Fig. 2E)**. Taking these results together, we conclude that a wide range of *cngc2* phenotypes, including enhanced pathogen resistance, depend on functional *YUC6*.

**Figure 2.**
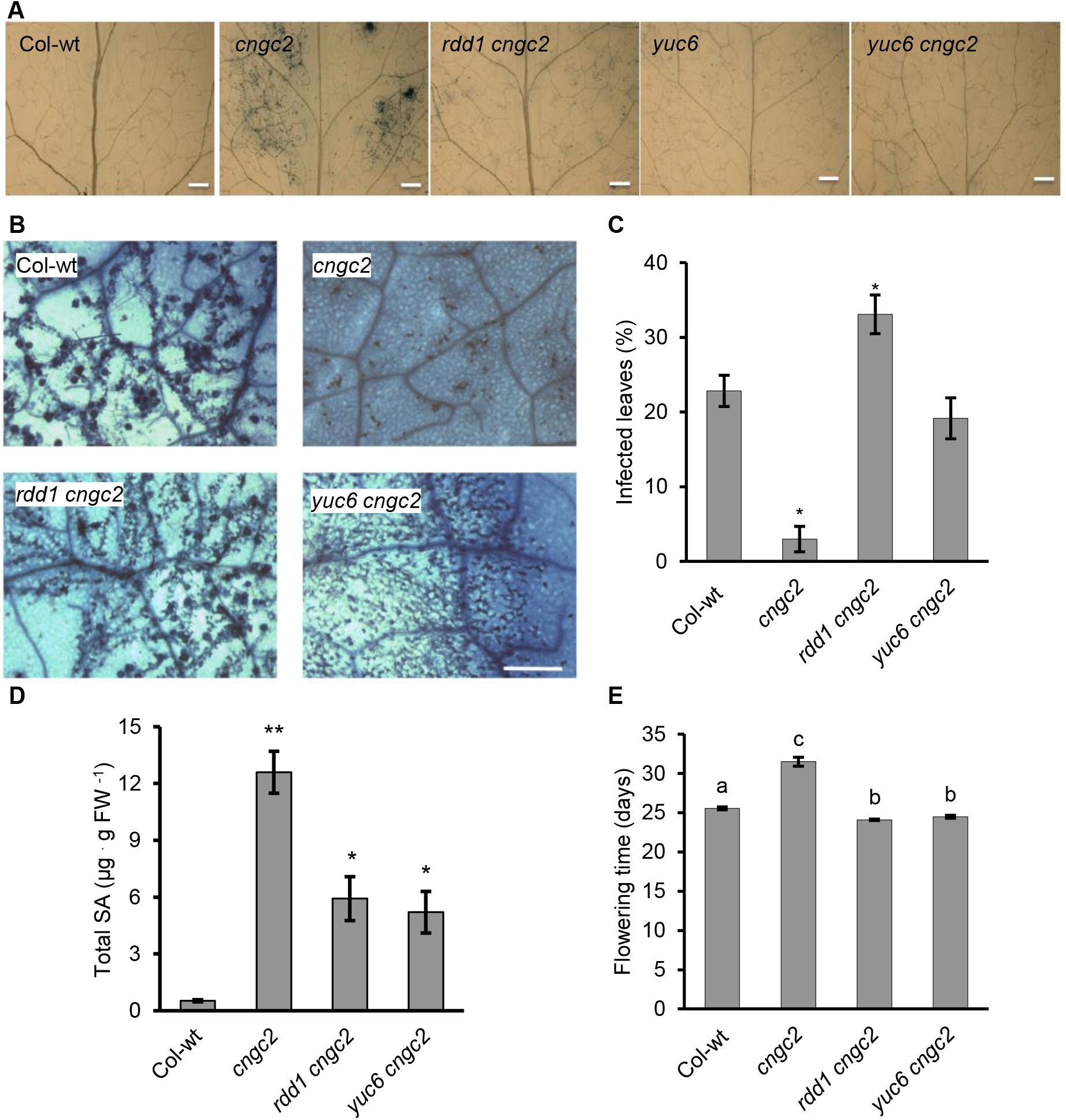
*yuc6* suppresses *cngc2*-conferred phenotypes. A, Trypan blue staining reveals a reduction in spontenous cell death in *yuc6 cngc2 and rdd1 cngc2* compared to *cngc2*. Scale bar = 0.5 mm. B, Breakdown of enhanced resistance phenotype of *cngc2* in *rdd1 cngc2* and *yuc6 cngc2* double mutants upon infection with *H. arabidopsidis* (Hpa), isolate Noco2. Trypan Blue staining of Col wild-type and mutants after inoculation with Hpa at a suspension of 8 × 10^5^ spores mL^-1^. Bar = 1 mm. C, Disease severity as a percentage of leaves showing symptoms of sporangiophore formation. Bars marked with an asterisk indicate significant difference from Col-wt (Student’s t-test, P < 0.05). n = 6. D, Total salicylic acid (SA) levels in 3-to 4-week-old wild-type and mutant leaves. SA levels are significantly different in *rdd1 cngc2* and *yuc6 cngc2* from *cngc2*. Error bars indicate standard error of the mean of 3 replicates. Bars marked with an asterisk indicate significant difference from Col-wt (Student’s t-test, P < 0.05). The experiment was repeated 3 times. E, *rdd1* and *yuc6* repress the delayed flowering phenotype of *cngc2. cngc2* plants exhibit delayed flowering compared to Col-wt, *rdd1 cngc2* and *yuc6 cngc2*. Shown are averages ± SE, n = 20-32. Bars marked with the same letter indicate no significant difference (Student’s t-test, P < 0.05).

### CNGC2 negatively affects TAA1/YUC6-mediated auxin biosynthesis

Since *rdd1* is a loss-of-function allele of *YUC6*, we postulated that auxin homeostasis is mis-regulated in *cngc2* mutants. In line with our expectations, *cngc2* contained more endogenous IAA than the WT control, in shoots and roots in mature plants, and this was partially reversed in the *rdd1 cngc2* and *yuc6 cngc2* double mutants (**Fig. 3**). This observation was statistically significant in shoots, but not in roots; however, the same trend has been observed in roots repeatedly. Thus, we think the difference between shoot and root samples is due to technical difficulties to collect mature plant root samples. This corroborates the enhanced dwarfism of *yuc6-1D cngc2* plants as *yuc6-1D* also hyper-accumulates auxin **(Fig. 1)**. These results are consistent with CNGC2 negatively affecting YUC6-dependent auxin biosynthesis/homeostasis.

**Figure 3.**
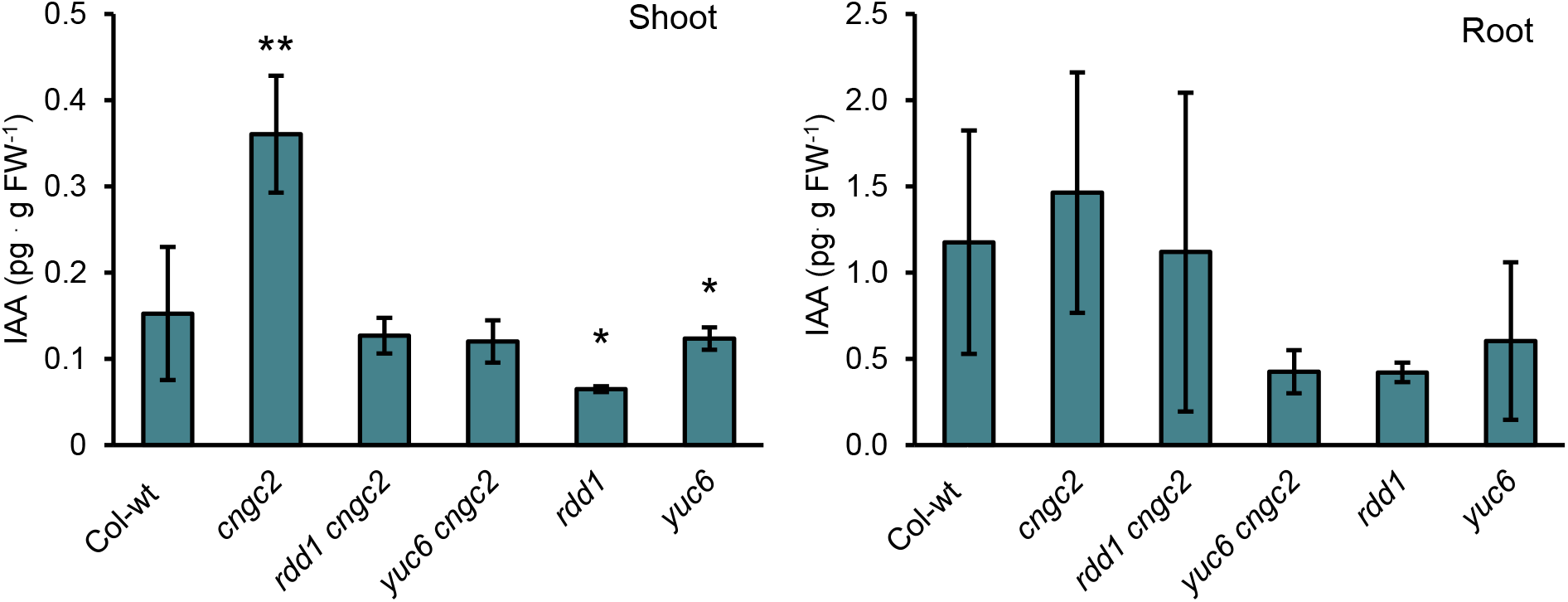
CNGC2 negatively affects TAA1/YUC6-mediated auxin biosynthesis. Shoot and root indole-3-acetic acid (IAA) levels of 5-week-old plants were measured using LC-MS/MS. The experiment was repeated 4 times and averages from one representative trial are presented; shown are ± SE, n = 3. Bars marked with an asterisk indicate significant difference from Col-wt (Student’s t-test, P < 0.05).

To discriminate between a YUC6-specific effect and a general, auxin biosynthesis-related effect, we assessed the requirement for another auxin biosynthetic gene, *TAA1*. The *TAA1* loss-of-function allele *weak ethylene insensitive 8* (*wei8-1*) exhibits gravitropic defects and its allelic mutant of *CK-INDUCED ROOT CURLING 1* (*ckrc1*) additionally exhibits reduced IAA levels, due to defects in the production of the substrate of YUCCAs for IAA biosynthesis (Mashiguchi et al., 2011; Stepanova et al., 2008; Zhou et al., 2011). We generated *wei8-1 cngc2* double mutants to assess the dependence of *cngc2*-associated phenotypes on IAA biosynthesis. These double mutants exhibited a similar suppression of *cngc2* phenotypes as *rdd1 cngc2* (**Fig. 4A**) and suppressed the elevated SA levels of *cngc2* **(Fig. 4B)**. Similar to *rdd1, wei8-1* also partially rescued the delayed flowering of *cngc2* **(Fig. 4C, D)**. Taken together, these data support the hypothesis that a wide variety of *cngc2* phenotypes depend on hyperactive auxin biosynthesis, rather than YUC6 function itself.

**Figure 4.**
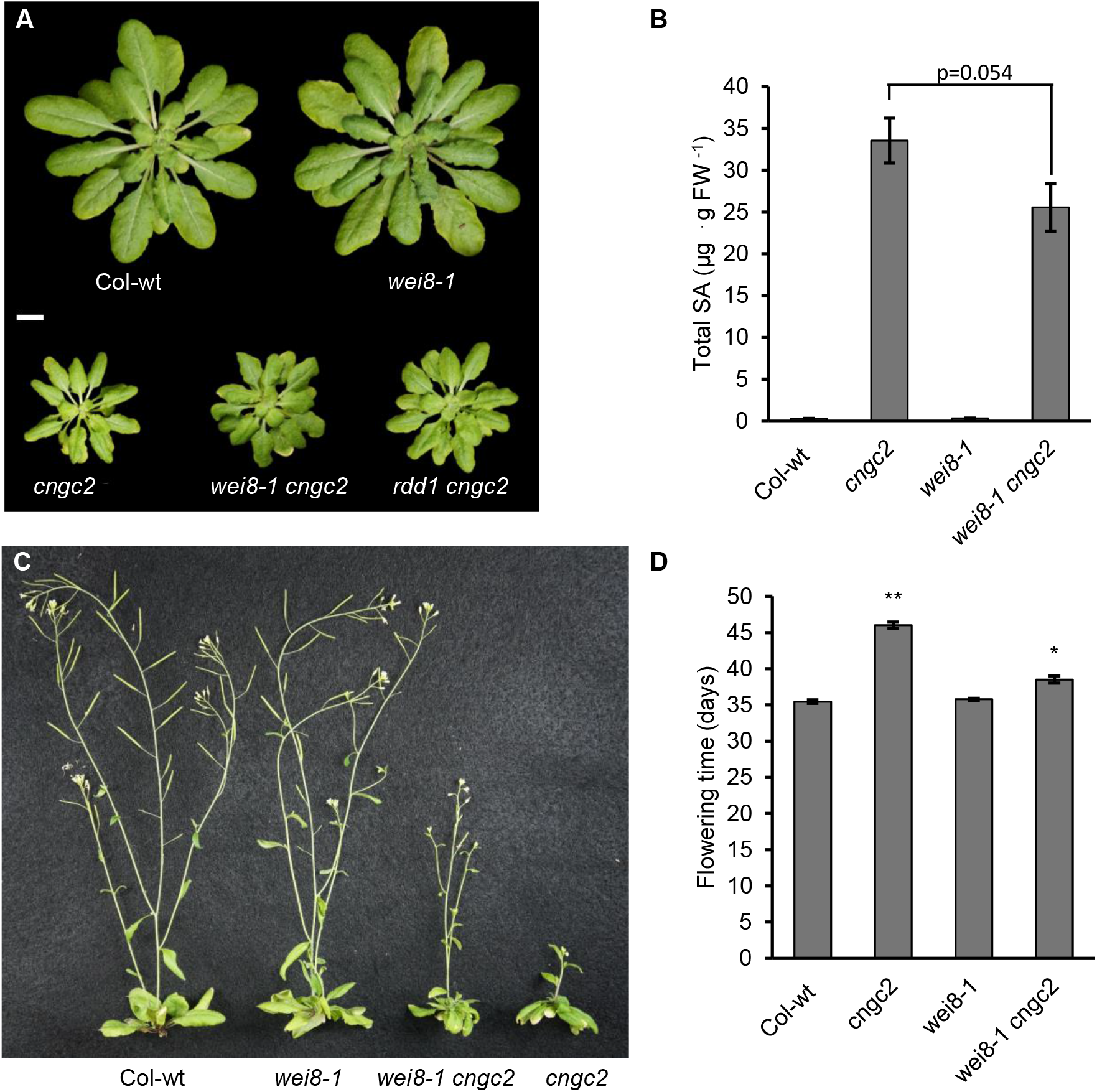
*wei8-1* (*taa1*) partially suppresses *cngc2*-conferred phenotypes. *A*, Morphology of 5-week-old Col-wt and mutant plants grown in short day conditions (9L:15D). *wei8-1 cngc2* partially suppress *cngc2*-conferred dwarf morphology to the same degree as *rdd1 cngc2*. Scale bar = 1 cm B, Total salicylic acid (SA) levels in 5-to 6-week-old Col-wt and mutant leaves. Error bars indicate standard error of the mean of 3 replicates. C, *wei8-1* represses the delayed flowering phenotype of *cngc2*. Picture was taken at 48 days D, Time to flowering is partially delayed in *wei8-1 cngc2*. Flowering time was measured in Col-wt and mutants by determining the emergence of the first bud. *cngc2* plants exhibit delayed flowering compared to Col-wt and *wei8-1*. Shown are averages ± SE, n = 23. Bars marked with an asterisk indicate a significant difference from Col-wt (Student’s t-test, P < 0.05).

### RDD1 antagonizes CNGC2-dependent auxin signaling

The *rdd1* mutation suppresses almost all *cngc2* mutant phenotypes (Chin et al., 2013). Morphological defects of *cngc2* that differ from phenotypes of other SA-related mutants suggest alterations in hormones related to development, such as auxin. Thus, we investigated the auxin-related phenotypes of *cngc2* and the *rdd1 cngc2* double mutant.

We found a delayed gravitropic root tip bending response in *cngc2*, which was partially rescued in *rdd1 cngc2* especially at earlier time points before 10 hours (**Fig. 5A**). We then analyzed the expression of the canonical auxin signaling output reporter, DR5 fused to GUS (DR5::GUS), as well as the Aux/IAA-based TIR1/AFB-activity sensor, DII-VENUS (Brunoud et al., 2012; Ulmasov et al., 1997). Auxin treatment elicited a strong auxin-responsive induction of DR5::GUS activity in Col wt but not in *cngc2* roots **(Fig. 5B)**. This indicates that the transcriptional auxin response is impaired in *cngc2*. Importantly, the *rdd1* mutation nearly completely restored auxin responsiveness of DR5::GUS in *rdd1 cngc2* roots (**Fig. 5B)**, suggesting that RDD1 is involved in the control of auxin signaling by CNGC2.

**Figure 5.**
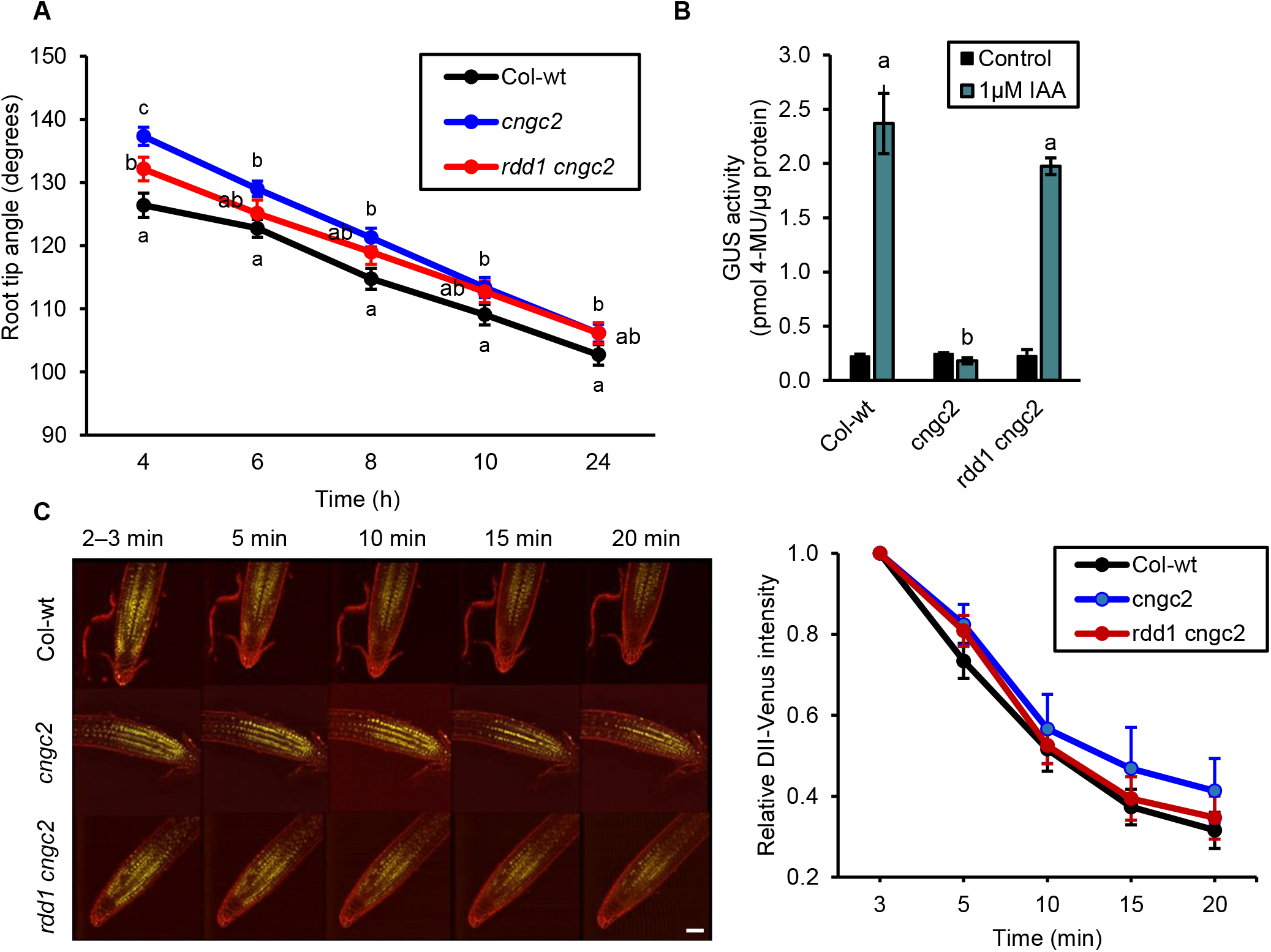
*cngc2* exhibits alterations in auxin-sensitivity, which is partially rescued in *rdd1 cngc2*. A, *cngc2* exhibits a delay in its gravitropic response. Time-course of gravitropic response of 7-day old seedlings measured every 2 hours from 4 hours till 24 hours after rotating the plate by 90°. *cngc2* displays delayed gravitropic root bending compared to wildtype (Col-wt) this is partially recovered in *rdd1 cngc2*. Shown are averages ± SE (n=70). Data points marked with the same letter indicate no significant difference (Student’s t-test, P < 0.05). B, GUS activity of DR5::GUS transgenic plants (Col-wt, *cngc2, rdd1 cngc2*). *cngc2* is insensitive to the application of 1µM exogenous IAA to its roots. This insensitivity is completely rescued in *rdd1 cngc2* root. Shown are averages ± SE, n = 3. Bars marked with the same letter indicate no significant difference (Student’s t-test, P < 0.05). C, DII:VENUS fluorescence of the primary root after the addition of 1µM IAA. (Left) Shown are representative images. (Right) Quantification of signal intensity of DII:VENUS at region of interest, relative to two minutes after the addition of 1µM IAA. Degradation of VENUS signal is delayed in *cngc2* relative to Col-wt and *rdd1 cngc2*. Shown are averages ± SE, n = 5. All experiments were repeated 3 times with comparable results. Scale = 50µm.

Auxin activates DR5::GUS activity through the TIR1/AFB-dependent ubiquitination of Aux/IAA proteins and their subsequent proteolysis. Therefore, we monitored the activity of the TIR1/AFB co-receptor via the Aux/IAA degradation sensor DII-VENUS (Brunoud et al., 2012). Auxin treatment triggered rapid decay in the DII-VENUS signal over a 20-min period in 5-day old roots in the Col-wt. This decay was slower and dampened in the *cngc2* background and was partially rescued in *rdd1 cngc2* roots **(Fig. 5C)**. Collectively, these results suggest that functional CNGC2 is required for TIR1/AFB-mediated auxin signaling. Moreover, the consistent suppression of the *cngc2* auxin-insensitivity implicates *RDD1* as a negative regulator of CNGC2 signalling.

### CNGC2 is required for auxin-induced Ca^2+^ signaling in the root

A non-transcriptional branch of SCF^TIR1/AFB^-based auxin perception triggers Ca^2+^ entry into the cell in a CNGC14-dependent manner (Shih et al., 2015; Dindas et al., 2018). Given that CNGC2 is required for SCF^TIR1/AFB^ activity in the context of transcriptional auxin signaling, we predicted a defect in auxin-induced Ca^2+^ signaling in *cngc2*. Therefore, we analyzed the auxin-induced Ca^2+^ response in WT, *cngc2*, and the *rdd1 cngc2* double mutant.

The highly sensitive FRET-based Ca^2+^ sensor Yellow Cameleon (YC)-Nano65 (Choi et al., 2014) responded rapidly to auxin treatment at various regions of interest (ROI) along the root (**Fig. 6B**). Auxin treatment at the root tip of WT rapidly induced a strong peak in [Ca^2+^]_cyt_, followed by a more sustained Ca^2+^ response (ROI1, **Fig. 6A, C**). The peak response was less pronounced in more shootward ROIs (ROI2, ROI3, and ROI4), but was followed by a clear sustained Ca^2+^ response over at least 120 s. This observation in WT was in line with previous reports using other Ca^2+^ reporters (Monshausen et al., 2011; Shih et al., 2015; Waadt et al., 2017). However, the Ca^2+^ increase upon auxin treatment was much weaker in *cngc2*. This defect was largely recovered in *rdd1 cngc2* **(Fig. 6A, C)**. Application of cold water elicited identical Ca^2+^ signals in WT and *cngc2*, indicating that the impairment in IAA-induced Ca^2+^ signals in *cngc2* is not related to a general disruption of Ca^2+^ signaling **(Supplemental Fig. S4)**. Taken together, these data demonstrate a requirement for CNGC2 in auxin-mediated Ca^2+^ signaling that is dependent on RDD1.

**Figure 6.**
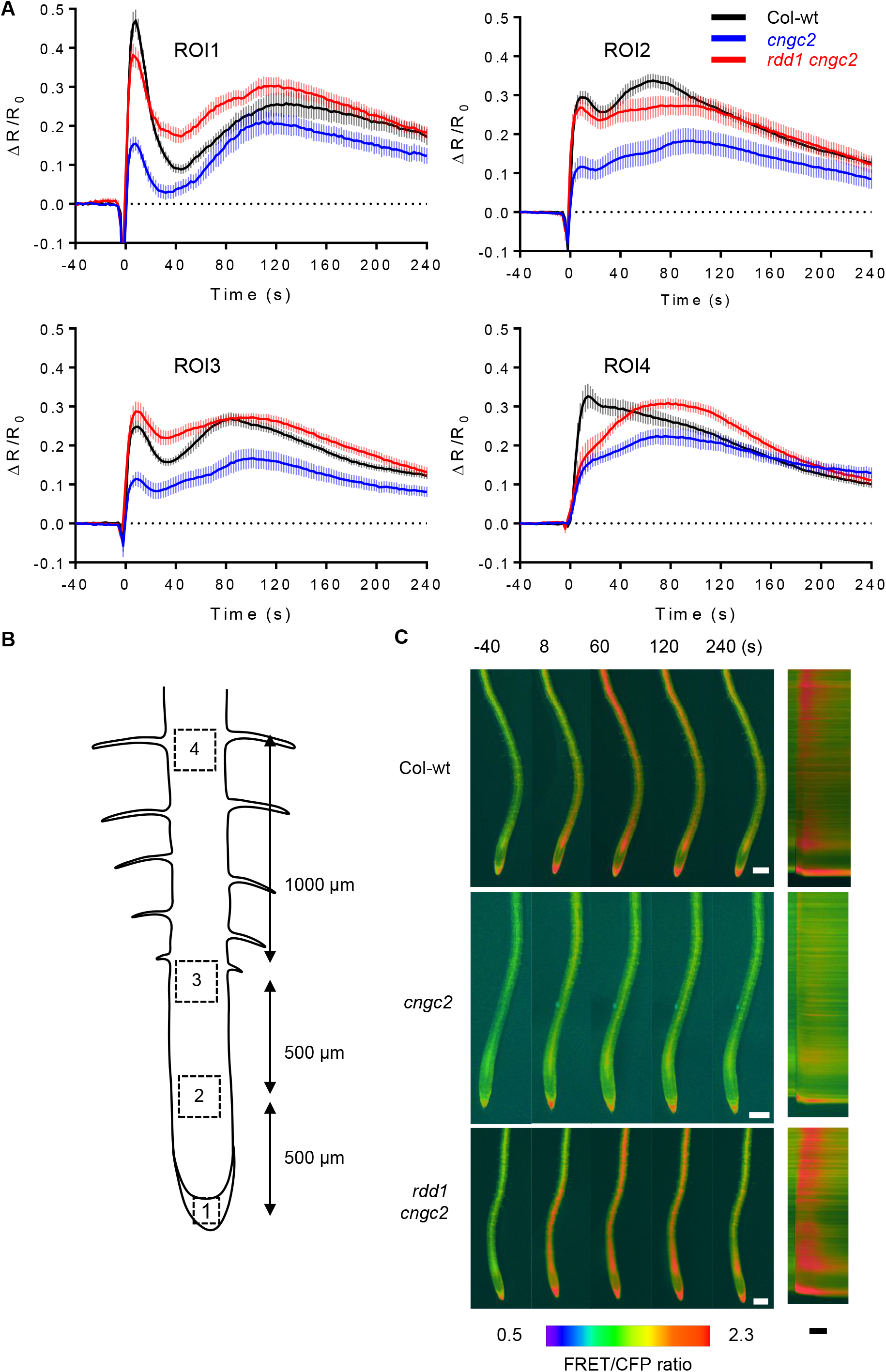
Auxin-mediated Ca^2+^ signals are defective in *cngc2* but rescued in *rdd1 cngc2*. A, Quantitative analysis using the FRET based Ca^2+^ sensor YCNano-65. The application of 1µM IAA at 0 s induces Ca^2+^ spikes in columella cells (ROI1), the elongation zone (ROI2 and ROI3) and the maturation zone (ROI4) of Col-wt root, but not in these corresponding regions of *cngc2*. This is largely recovered in *rdd1 cngc2* (n = 10-12). B, Schematic of region of interests (ROI) of the *A. thaliana* root that were examined for auxin-induced Ca^2+^ signals. 1 µM IAA was applied to the root tip region around ROI1. C, Kymograph analysis of changes in FRET/CFP ratio upon the addition of 1µM IAA in whole root of transgenic lines expressing yellow cameleon-Nano65 in Col-wt, *cngc2* and *rdd1 cngc2* background. White scale bar = 0.2 mm, black scale bar = 100 s. Movies are included as a supplemental data.

## DISCUSSION

In the two decades since the discovery of the autoimmunity phenotype of *CNGC2* loss-of function mutants, CNGC2 has been the most intensively studied plant CNGC. However, *cngc2* exhibits pleiotropic phenotypes, which are unique among conventional immunity mutants (Moeder et al., 2011, 2019), This indicates its wider physiological role beyond immunity and raises the fundamental question of whether a specific plant CNGC can generate different downstream signals depending on stimuli (Dietrich et al., 2020). To examine this question, we embarked on a suppressor screen of *cngc2* and discovered an unexpected, tight connection between CNGC2 and auxin biosynthesis and signaling.

### CNGC2 suppresses SCF^TIR1/AFB^-mediated auxin signaling

Auxin perception through the well-described SCF^TIR1/AFB^-based system activates transcription by proteolysis of Aux/IAA proteins (Leyser, 2018). Upon auxin treatment, the SCF^TIR1/AFB^ receptor also activates CNGC14 via an unknown mechanism (Dindas et al., 2018; Shih et al., 2015). Here, we show that CNGC2 is required for SCF^TIR1/AFB^ auxin signaling at the level of transcriptional regulation (DR5::GUS reporter analysis), as well as the induction of Ca^2+^ signals. This is reflected in the reduced root gravitropism observed in the *cngc2* mutant.

Since CNGCs were suggested to form heterotetrameric channels (Chin et al., 2013; Pan et al., 2019; Tian et al., 2019) and CNGC14 plays a role in auxin-related Ca^2+^ signaling (Shih et al.; 2015, Dindas et al., 2018), it is possible that CNGC2 and 14 form a functional Ca^2+^ channel conducting auxin-induced Ca^2+^ influxes. Indeed, both CNGC14 and CNGC2 localize at the plasma membrane and are required for auxin-induced Ca^2+^ influx into the cytosol from the extracellular apoplastic Ca^2+^ pool (Lemtiri-chlieh & Berkowitz, 2004; Wang et al., 2017; Dindas et al. 2018; Shih et al., 2015). Moreover, the gravitropic response defects of *cngc2* and *cngc14* were quite similar under our experimental conditions (**Supplemental Fig. S5A)**. However, in contrast to the pleiotropic developmental defects seen in *cngc2* (Clough et al., 2000), *cngc14* exhibits little to no additional developmental defects besides the reduced gravitropism and root hair phenotypes (Dindas et al., 2018; Shih et al., 2015; Brost et al., 2019). An additional obvious difference from *cngc2* mutants is the absence of defense-related phenotypes in *cngc14* (Brost et al., 2019; Dindas et al., 2018; Shih et al., 2015). These differences together with the limited overlap in expression patterns (**Supplemental Fig. S5B**) indicate that CNGC14 and CNGC2 likely have different biological functions and argues against CNGC2 being a component of the SCF^TIR1/AFB^-CNGC14 module and/or forming a heterotetrameric channel with CNGC14. Therefore, we favor a scenario in which CNGC2-mediated Ca^2+^ signals control auxin biosynthesis and the activity of SCF^TIR1/AFB^ -auxin sensing independent from CNGC14. This also provides a straightforward explanation for the reduced auxin sensitivity of DR5::GUS and DII-VENUS auxin reporters in *cngc2*.

### CNCC2-CNGC4 mediated Ca^2+^ signaling suppresses auxin biosynthesis

Whether CNGC2 can form a heterotetramer with CNGC14 remains to be seen; however, CNGC2 is known to form a functional Ca^2+^ channel through heteromerization with CNGC4 (Chin et al., 2013, Tian et al., 2019). The respective mutants, *cngc2* (*dnd1*) and *cngc4 (hlm1*/*dnd2*) have identical morphological and molecular phenotypes. To date, these phenotypes have mainly been characterized with respect to their autoimmunity phenotypes (Clough et al., 2000, Balagué et al., 2003), and were recently shown to be defective in PAMP-induced Ca^2+^ signals (Tian et al., 2019). Here, we found that *cngc2* exhibits hyper-accumulation of endogenous IAA similar to *cngc4* (Kale et al., 2019). This observation indicates that a CNGC2–CNGC4 heteromeric channel generates a Ca^2+^ signal that suppresses IAA biosynthesis. The auxin hyperaccumulation in *cngc2*, and most of its pleiotropic phenotypes, could be repressed when the auxin biosynthesis gene *YUC6* was mutated. Moreover, the phenotypes of *cngc4* and even the double mutant *cngc2 cngc4* could also be suppressed by the *yuc6* allele *rdd1* (Chin et al., 2013), suggesting that our findings about auxin signaling and biosynthesis in *cngc2* can be extrapolated to *cngc4*.

In addition to its FMO function for IAA biosynthesis, YUC6 also exhibits thiol reductase activity, which plays a role in ROS homeostasis (Cha et al., 2016; Cha et al., 2015). Since hyper-accumulation of ROS is a common phenomenon associated with many lesion mimic mutants, including *cngc2*, it is possible that the thiol reductase activity rather than the FMO function of YUC6 is related to the suppression of *cngc2* phenotypes. However, the *rdd1* mutation is not located in the thiol reductase domain (Fig. 1A). Furthermore, loss-of-function of TAA1/WEI8, which functions one step upstream of YUC6 in the IAA biosynthesis pathway, also rescues *cngc2* phenotypes. This strongly suggests that alterations in IAA biosynthesis, not the thiol reductase activity of YUC6, lead to the suppression of *cngc2*. The *taa1* (*wei8-1*) mutation had a slightly weaker effect on *cngc2* than *yuc6*, probably due to redundancies with TRYPTOPHAN AMINOTRANSFERASE RELATED PROTEIN 1 and 2 (TAR1, TAA2) (Stepanova et al., 2008). Taken together, the unique morphological and physiological phenotypes of *cngc2* and *cngc4* among lesion mimic mutants can be explained by a hyperactive TAA–YUC auxin biosynthesis pathway. Indeed, other lesion mimic mutants such as *suppressor of npr1-1, constitutive1* (*snc1)* and *constitutive expressor of PR genes 6* (*cpr6)* exhibit lower levels of endogenous IAA (Wang et al., 2007), further supporting this notion. Recently, auxin perception via the TRANS-MEMBRANE KINASE 4 (TMK4) receptor-like kinase, was shown to suppress auxin biosynthesis via phosphorylation of TAA1 (Wang et al., 2020). Therefore, it will be interesting to see if TMK4 acts together with CNGC2 to suppress auxin biosynthesis.

### CNGC2-mediated Ca^2+^ signals at the nexus of immunity and development

As mentioned, the autoimmunity mutants *cngc2* and *cngc4* have almost identical phenotypes including elevated SA levels, which are suppressed by *rdd1* (Chin et al., 2013). Thus, the identification of *RDD1* as the auxin biosynthesis gene *YUC6* initially led us to hypothesize that the effect of *rdd1* is simply restoring the balance in the SA–auxin antagonism in *cngc2*. Auxin treatment promotes disease symptoms and prevents the full induction of the SA-inducible antimicrobial gene *PATHOGENESIS RELATED1* (*PR1)*, indicating that auxin antagonizes defense responses via suppression of SA signaling. Indeed, many pathogenic microorganisms manipulate plant immunity through modification of auxin signaling by their effectors and/or auxin originating from the microorganisms themselves (Kazan and Lyons, 2014; Kunkel and Harper, 2018; Mutka et al., 2013). Moreover, treatment with SA or the SA analog BTH globally suppresses transcription of auxin-related genes and various autoimmune mutants exhibit lower IAA levels and reduced sensitivity to auxin (Wang et al., 2007). However, the *cpr22* autoimmune phenotype, caused by the *CNGC11*/*12* gain-of-function mutation, which induces constitutive activation of Ca^2+^ signals (Moeder et al., 2019), was not suppressed by *rdd1* (Fortuna et al., 2015). Furthermore, blocking SA biosynthesis by introducing the *sid2* mutation reverted almost all phenotypes of *cpr22* (Yoshioka et al., 2006), while the *sid2 cngc2* and *sid2 cngc4* double mutants retain clear morphological defects (Genger et al., 2008). Thus, simple SA–auxin antagonism is not the cause of the suppression of *cngc2* and *cngc4* phenotypes by *rdd1*.

These data also suggested that CNGC2/CNGC4 activate autoimmunity through distinct pathways, indicating a unique aspect in CNGC2/CNGC4-mediated immunity among lesion mimic mutants. Thus, *CNGC2* may primarily play a role in auxin homeostasis/signaling and its observed immunity phenotypes (i.e., SA accumulation) are a consequence of this auxin-related defect. Alternatively, *CNGC2* may act independently or at a pivotal point in multiple physiological processes, such as defense and development. Indeed, *CNGC2* has recently been implicated in the regulation of SAM size through the CLAVATA signaling cascade and Ca^2+^ homeostasis (Chou et al., 2016; Wang et al., 2017) as well as PAMP-induced immunity (Tian et al., 2019). Ca^2+^ is a universal secondary messenger and is involved in almost all aspects of cellular signaling. It is possible that CNGC2 generates distinct Ca^2+^ signals (or Ca^2+^ signatures) depending on the stimulus and acts at the nexus of immunity and development. Such specificity could be achieved through changing channel subunit composition or by being part of a channelosome with stimuli-specific signaling components, such as receptors and decoders (Dietrich et al., 2020).

Taking our results together, we propose that plasma membrane localized CNGC2 forms a heterotetrametric channel with CNGC4 and generates Ca^2+^ signals that affect the TAA–YUC auxin biosynthetic pathway, likely to prevent over accumulation of IAA. CNGC4 has been reported to form a heterotetramer with CNGC2 (Chin et al., 2013, Tian et al., 2019) and *cngc4* also exhibits abnormal accumulation of IAA (Kale et a., 2019). Since the loss-of-function mutants of CNGC2 are impaired in their ability to generate a Ca^2+^ influx upon IAA treatment, the regulation of the TAA–YUC auxin biosynthetic pathway by CNGC2/CNGC4 must be a feedback regulation for auxin homeostasis, similar to that proposed by Wang et al. (2020) for TMK4. In addition, CNGC2 function is required for TIR/AFB-mediated auxin signaling; thus, CNGC2/4 mediated Ca^2+^ signals may control the activity of SCF^TIR1/AFB-^auxin sensing directly, or via facilitating proper IAA homeostasis. In this scenario, *cngc2* has abnormal accumulation of IAA resulting in the desensitization of cellular auxin signaling and morphological defects as a long-term effect. Alternatively, Ca^2+^ signals generated by CNGC2 may directly affect auxin signaling in addition to its biosynthesis. In any case, CNGC2 must play a role immediately after perception of auxin or associated to the perception itself. Although at this point, we cannot exclude other possibilities such as indirect effects, this model can be one plausible scenario. Further investigation of direct downstream targets of CNGC2-mediated Ca^2+^ influx will be necessary.

For two decades, CNGC2 has been studied intensively from an immunity point-of-view. However, publications in recent years as well as our current work strongly indicate a more diverse range of functions for CNGC2. Thus, the current study contributes to a more comprehensive understanding of CNGC2-mediated Ca^2+^ signaling and reveals the importance of CNGC2 beyond its role in defense.

## MATERIALS AND METHODS

### Plant Materials and Growth Conditions

For phenotypic analyses, *Arabidopsis thaliana* seeds were cold stratified at 4°C for two days prior to being grown at 23°C on Sunshine-Mix #1 (Sun Gro Horticulture, Vancouver, Canada). Plants were grown in a growth chamber with a 9-h photoperiod (9-h light/ 15-h dark) and a day/night temperature regime of 22°C/18°C. This condition prolongs the growth stage and induces clearer morphological phenotypes of the mutants. For other experiments, plants were grown on Petri dishes with half-strength Murashige and Skoog (½ MS) medium, 1% sucrose, and 0.8% (w/v) agar at pH 5.8 under ambient light conditions. Sterile media was either supplemented with IAA of the desired concentration or equal volumes of ethanol as a solvent control. Homozygous mutants were identified by PCR using primers listed in **Supplementary Table S4**. Since *rdd1* was isolated using the T-DNA insertion allele *cngc2-3* (Chin et al., 2013), we have used *cngc2-3* throughout this work except for the Ca2+ imaging analysis (see FRET-based Ca2+ analysis using YC-Nano65 section).

### Identification of *rdd1* as an allele of *YUC6*

Illumina HiSeq 1500 system (Illumina, Inc., San Diego, CA, USA) at the McMaster Institute for Molecular Biology and Biotechnology (MOBIX) was used to sequence the genomic DNA extracted from leaves of 4-week old *rdd1 cngc2-3* and *cngc2-3* plants. DNA was extracted from 100 mg of powdered leaf tissue using the CTAB method (Thompson, 1980) with a slight modification: 200 units/ml of RNAse A and 0.4 g/ml PVP (MW 40000; Sigma-Aldrich, Canada) were added to the CTAB extraction buffer. Extracted DNA was precipitated twice with isopropanol, once with 75% EtOH and resuspended in 0.1X TE buffer supplemented with 0.2 units of RNAse A. DNA integrity was assessed using agarose gel electrophoresis and quantified with a Quant-iT PicoGreen dsDNA Assay Kit (Thermo Fisher Scientific) on a Bio-Tek Powerwave HT microplate reader. Multiplex libraries were prepared according to the manufacturer’s instructions (Nextera) and the three libraries were sequenced on one flow-cell lane in high throughput mode. Raw sequence reads were trimmed for adaptor sequences with Cutadapt software (Martin, 2011) and unpaired reads, and those of length <36 nt, were excluded from further analyses. Paired reads were aligned against the TAIR10 WT reference genome using BWA (Li & Durbin, 2009), and sequence variants were detected with the mpileup function of SAMtools (Li et al., 2009).

### GUS enzymatic assay

Seedlings were grown on ½ MS, 1% sucrose, and 0.8% (w/v) agar plates for six days before being transferred to plates supplemented with 1 µM IAA, or solvent alone (ethanol at final concentration of 0.09%), for 24 hours. GUS reporter activity was analyzed at an excitation wavelength of 365 nm and emission wavelength of 455 nm, using a TECAN plate reader (every 10 min for 2 h) in the presence of 4-methylumbelliferyl glucuronide (4-MUG). GUS activity was standardized against protein concentration and data was reported as GUS activity in pmol 4-methylumbelliferone (4MU) per µg protein.

### Pathogen infection

Infection with *Hyaloperonospora arabidopsidis* isolate Noco, which is virulent to the WT accession of *Arabidopsis* was performed as described previously with 5 × 10^5^ spores per ml (Chin et al., 2013).

### Analysis of endogenous salicylic acid

Endogenous SA was analyzed using the *Acinetobacter* sp. ADPWH lux-based biosensor as previously described (Defraia et al., 2008).

### Gravitropic root bending

Five to seven-day-old seedlings were photographed every two hours between 4 and 10 hours, and then again at 24 hours, from the start of gravistimulation. The deviation in root tip angle from 90° was analyzed using the angle tool on ImageJ software (http://rsbweb.nih.gov/ij/).

### Analysis of flowering transition time

*Arabidopsis thaliana* wildtype and mutant plants were grown on Sunshine-Mix #1 (Sun Gro Horticulture, Vancouver, Canada) in a growth chambers under 16-h photoperiod (16-h light/ 8-h dark) at 22°C/18°C. Observations were made every other day and floral transition was recorded as days taken for first bolt to form from time of sowing as described in Fortuna et al., 2015.

### DII-VENUS analysis

For DII-VENUS analysis, seedlings were visualized for 20 minutes immediately after supplementation with 1 µM IAA. Confocal images were captured using a Leica TCS SP5 confocal system with an acousto-optical beam splitter (HCX PL APO CS 40x immersion oil objective; numerical aperture, 1.25), and the acousto-optic tunable filter 514 for the argon laser using the nm output, set at 20%. The detection window was set to 525 to 600 nm for YFP (Leica Microsystems). Seven-to nine-day-old seedings were stained with 1x propidium iodide and imaged at 40X magnification. Images were processed using Leica Las AF lite software.

### FRET-based Ca^2+^ analysis using YC-Nano65

Multiple lines of Wt, *dnd1*, and *rdd1 dnd1* carrying YC-Nano65 were generated as previously described (Choi et al., 2014). These plants were grown on the surface of a vertical agar plate with ½ MS, 1% (w/v) sucrose, and 0.5% gellan gum at pH 5.8 under ambient light conditions. The root tips were treated with 10 μl of 1 μM IAA. FRET (cpVenus) and CFP signals from YC-Nano65 were acquired using a motorized fluorescence stereo microscope (SMZ-25, Nikon) equipped with image splitting optics (W-VIEW GEMINI, Hamamatsu Photonics) and a sCMOS camera (ORCA-Flash4.0 V2, Hamamatsu Photonics) as previously described (Lenglet et al., 2017; Toyota et al., 2018). For high-resolution confocal Ca^2+^ analysis, the transgenic plants were grown vertically under a thin layer (approximately 2 mm) of the growth medium (½ MS, 1% (w/v) sucrose and 0.5% gellan gum at pH 5.8) on a cover glass (24 × 50 mm, Fisher Scientific) for six days at 23°C. The tip of the root was exposed by removing a small window (approximately 500 μm × 500 μm) from the gel, and 10 μl of 1 μM IAA was applied to the root tip area through this window. FRET (cpVenus) and CFP signals from YC-Nano65 were acquired using a laser scanning confocal microscope (LSM780/Elyra; Newcomb Imaging Center, Department of Botany, University of Wisconsin, Madison) as previously described (Choi et al., 2014). The cpVenus/CFP ratio was calculated using 6D imaging and Ratio & FRET plug-in modules, and the kymograph of the entire root was generated over 240 seconds (NIS-Elements AR, Nikon).

### Accession Numbers

Sequence data from this article can be found in the EMBL/GenBank data libraries under accession number(s): *CNGC2* (AT5G15410), *RDD1/YUC6* (AT5G25620), *TAA1* (AT1G70560).

## Acknowledgements

We thank Dr. Catherine Chan for providing the Ws *cngc2* seeds, Drs. Barbara Kunkel and Yunde Zhao for providing the *Yucca6-1D* seeds, and Drs. Thomas Berleth and Enrico Scarpella for providing the DII-VENUS and DR5::GUS transgenic seeds. We appreciate the fruitful discussion and help from Dr. Matyáš Fendrych. Genomic sequencing was done with the help of Dr. Elizabeth Weretilnyk and the McMaster Institute for Molecular Biology and Biotechnology (MOBIX). We thank Drs. Andrew S Whiteley and Zhonglin Mou for providing the ADPWH lux-based salicylic acid biosensor. This work was supported by a Discovery grant from the National Science and Engineering Research Council (NSERC) to K.Y., a graduate student fellowship from NSERC to S.C., and KAKENHI (17H05007, 18H05491) to M.T.

## Author contributions

K.Y. and W.M. conceived the project; K.Y., W.M., S.V., T.B., M.T., S.G., E.N. designed the project; S.C., K.C., A.F., M.C., S.V., M.T., W.M. and E.N. performed the experiments and analyzed the data; S.C., K.Y., W.M. and S.V. analyzed data and wrote the article.

